# Fluorogenic DNase Sensor Reveals Ubiquitous DNase Activity in Podosomes and Invadopodia

**DOI:** 10.1101/2020.06.01.127902

**Authors:** Kaushik Pal, Yuanchang Zhao, Yongliang Wang, Xuefeng Wang

**Author notes:** These authors contributed equally to this work (Kaushik Pal, Yuanchang Zhao).

## Abstract

Podosomes and invadopodia, collectively termed invadosomes, are important adhesive and degradative units formed in macrophages, osteoclasts, dendritic cells, cancer cells, and many other cell types. Invadosomes are well known for recruiting proteases that degrade matrix proteins and facilitate cell invasion. In contrast to the extensively studied proteases, another important class of degradative enzymes, DNase, remains uninvestigated and in fact, unknown in invadosomes. Using surface nuclease sensor (SNS), which reports deoxyribonuclease (DNase) activity on the cell membrane by fluorescence signal, we revealed that invadosomes, regardless of cell types or species, universally recruit DNase and readily degrade extracellular double-stranded DNA (dsDNA). We identified the recruited DNase as GPI-anchored membrane protein DNase X which functions locally at the cell-substrate interface and is co-localized with the actin cores of the invadosomes. DNase recruitment is highly consistent and rapid in invadosomes. Co-imaging of F-actin and DNase activity shows that 46-86% invadosomes (dependent on cell types) have associated DNase activities. Time series imaging shows that DNase becomes active within a minute after the actin nucleation, functioning concomitantly with protease activity in podosomes but preceding it in invadopodia. Overall, this discovery suggests a richer arsenal of degradative enzymes in invadosomes at the cell-substrate interface. This work would likely prompt more studies to investigate DNase in invadosomes, in particular, to understand the physiological role of invadosome-associated membrane DNase in cell functions such as immune response, cell migration, matrix remodeling, etc.

## Introduction

Podosomes and invadopodia are micron-sized membrane protrusive structures that have both adhesive and degradative functions by interacting with the surrounding extracellular matrix (ECM)^1-2^. They are found in a wide range of cell types, with podosomes formed in monocytes, osteoclasts, smooth muscle cells, *etc*., and invadopodia formed in many carcinoma cells. Although podosomes and invadopodia are found in different cell types, and their temporal dynamics are quite different, they share a significant structural and enzymatic similarities^3^. Both structures have actin cores surrounded by peripheral adhesion rings consisting of integrin, vinculin, paxillin, etc.^4-5^. In recent literature, these two structures were often collectively termed invadosomes^6-7^. As formed in various cell types, invadosomes are highly versatile and involved in many physiological processes such as cell motility, cancer invasion, and metastasis, ECM remodeling, bone structure maintenance, *etc*^8-12^, therefore attracting significant research interest.

One characteristic feature of invadosomes is their enzymatically degradative function. Matrix degradation by invadopodia has been correlated with cancer metastasis and invasion^13-15^. In parallel, the degradative nature of podosomes has been considered as an important factor regulating the migration and tissue infiltration of macrophages^16-18^, the phagocytes specialized in removing cellular debris and pathogens. Due to the physiological and pathological importance of their enzymatic functions, invadosomes have been pursued as pharmaceutical targets^19^. Currently, the degradative ability of invadosomes is solely attributed to proteases such as serine proteases, ADAM proteases, and metalloproteases^20-23^. Surprisingly, DNase, another major class of enzymes found in the living system, has not been studied, identified, or even aware of in invadosomes.

In this work, we revealed that DNase is consistently and ubiquitously recruited to both podosomes and invadopodia. Previously, we developed a surface-immobilized nuclease sensor (SNS) that detects and images DNase activity at the cell-substrate interface in real-time by fluorescence microscopy^24^. Here, we tested both invadopodia (in human and mouse cancer cell lines) and podosomes (in human and mouse macrophage-like cell lines) on the SNS platform. Without exception, strong localized DNase activity was observed in the invadosomes. Spatio-temporal dynamics of structures and enzymatic activities monitored in real-time confirmed that DNase is recruited quickly after the actin core nucleation and remains active at the core region. We further identified the DNase in invadosomes as DNase X^25^, a glycosylphosphatidylinositol (GPI)-anchored membrane protein. We then devised a co-degradation assay, by which we revealed that DNase and proteases are active concomitantly in podosomes, whereas DNase activity preceded protease activity in invadopodia. These results suggest that podosomes are equipped with more active enzymes than known earlier. To our knowledge, this is the first report of DNase activity at the cell-substrate interface in invadosomes, and we believe this finding will be an important starting point to investigate more physiological relevance of DNase in invadosomes.

## Results and Discussion

### SNS selectively reports DNase in invadosomes

Previous studies have discovered a plethora of adhesive, structural, and enzymatic proteins in invadosomes (Fig. 1a)^26-27^. Many proteases are known to take active roles in the function of invadosomes. DNase, as another major class of degradative enzymes, somehow remains uninvestigated in invadosomes. Here we applied two DNase sensors and total internal reflection fluorescence (TIRF) microscopy to investigate the DNase activity in invadosomes at the cell-surface interface in both live and fixed cells. The first DNase sensor is surface-immobilized Cy3-labeled double-stranded DNA (dsDNA), as shown in Fig. 1b. This sensor loses the dye if degraded by DNase and therefore reports DNase activity by fluorescence loss, resembling the classic dye-labeled gelatin degradation assay used to report protease activity of invadopodia^28-29^. The other sensor is SNS (Fig. 1c), a dsDNA labeled by a quencher-dye pair at the opposite strand, and a biotin tag for surface immobilization through biotin-neutravidin interaction. Initially dark SNS becomes fluorescent if the dsDNA is degraded and dye is freed from the quencher, hence reporting DNase activity at the cell-substrate interface by fluorescence gain. Human macrophage-like cell line THP-1 (activated with TGF-*β*1^30^) were plated on glass surfaces coated with these two types of sensors, respectively. On the surface coated with dye-labeled dsDNA, strong fluorescence loss in a punctate pattern was observed, co-localizing with podosomes that were identified by the actin cores and vinculin rings (Fig. 1d, Figs. S1a, c). To rule out the possibility that these dark puncta were caused by local dye bleaching or sensor detachment, activated THP-1 cells were plated on an SNS-coated surface where podosomes produced positive fluorescence puncta, confirming that the DNA construct in the SNS was degraded (Fig. 1e, Figs. S1b, d). These two assays confirmed that podosomes in THP-1 cells exhibit strong DNase activity in the core regions. Further verification was conducted to confirm that the DNase in podosome is dsDNA-specific. Results showed that the DNase in podosomes degraded dsDNA regardless of DNA sequences, but did not degrade single-stranded DNA (ssDNA) or DNA/PNA (peptide nucleic acid) hybrid duplex which is supposed to be DNase-resistant (Fig. S2). Because the SNS sensor specifically responds to DNase activity, not dye bleaching, we adopted it for all following experiments. We evaluated the consistency of DNase activity in podosomes. In the co-imaging of F-actin and DNase activity of THP-1 cells, 86±5 % actin puncta are co-localized with SNS punctate signal (analyzed on the basis of 600 podosomes over 20 cells). The experiment was conducted more than three times, and the percentage did not vary significantly. Note that the actual percentage of invadosomes exhibiting DNase activity should be even higher than observed because DNase activity starts to appear ∼1 minute later after the actin core nucleation (see the study of temporal dynamics of DNase activity) and therefore a small set of actin puncta have yet to produce any SNS signal. In the images, many SNS puncta have no corresponding actin cores. This is because invadosomes are subject to disassembly, while SNS surface records all historic DNase activity. As a result, there is a subset of SNS puncta where invadosomes have already disassembled.

**Fig. 1.**
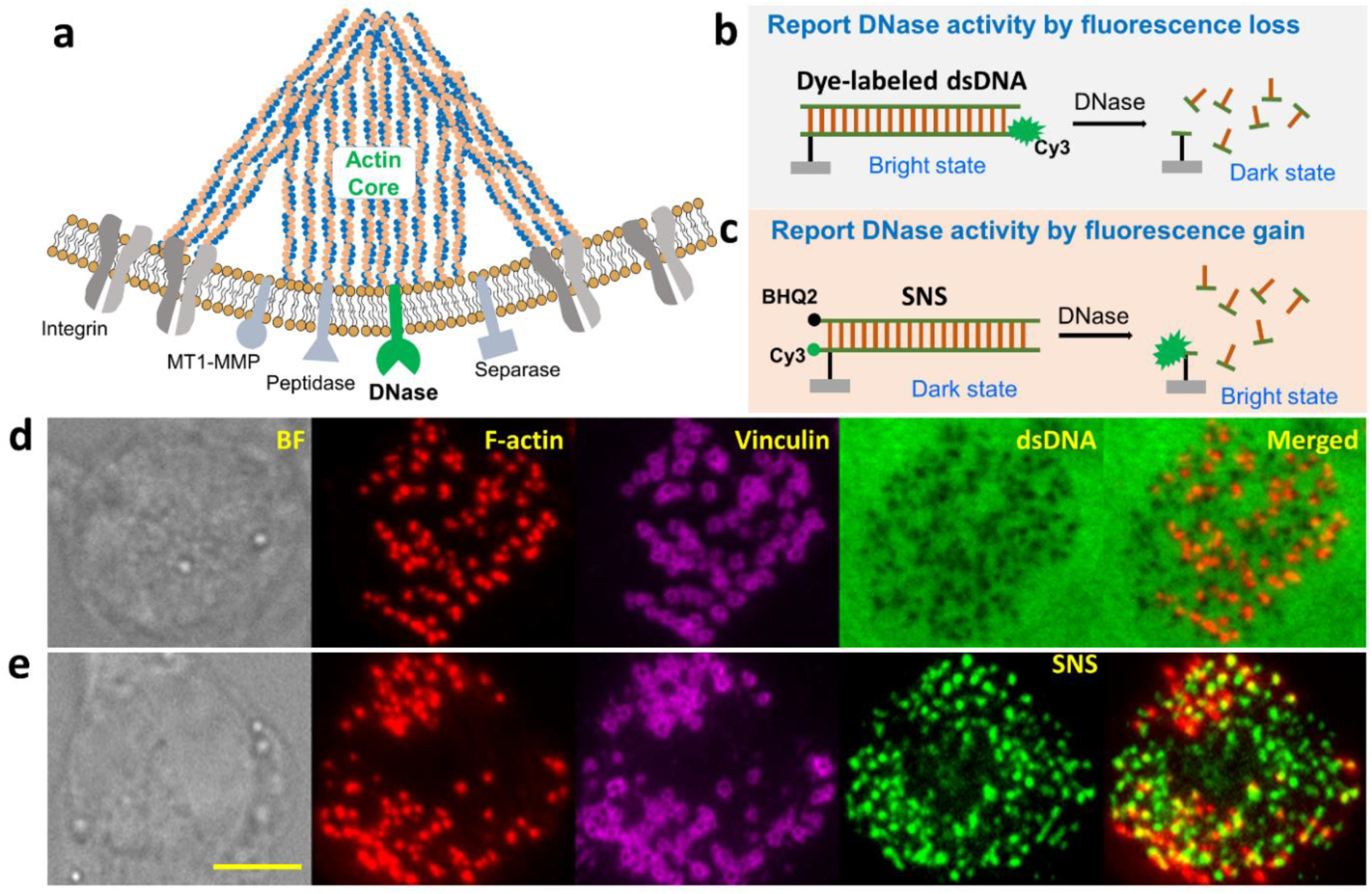
DNase activity in invadosomes reported by surface nuclease sensors (SNS). (a) A basic schematics of the invadosome structure. DNase depicted in green is identified in invadosomes in this study. (b) Surface-immobilized Cy3-labeled dsDNA visualizes DNase activity by fluorescence loss in response to DNA degradation caused by local DNase. (c) SNS with a quencher-dye pair visualizes DNase activity by fluorescence gain, as the degradation of DNA frees the Cy3 from quenching. (d) A THP-1 cell on a surface coated with Cy3-labeled dsDNA. Podosomes were identified by the actin core and vinculin ring. dsDNA was degraded in a dark punctate pattern co-localized with podosomes. (e) A THP-1 cell on an SNS immobilized surface, which showed DNA degradation in a bright punctate pattern co-localized with podosomes. Statistics show that 86±5% actin puncta are co-localized with SNS punctate signal (600 podosomes over 20 cells). Scale bar: 5 µm.

### DNase is ubiquitously recruited to invadosomes

To test if DNase is commonly recruited to invadosomes, we have applied SNS assay to four cell types from different species, which form either podosomes or invadopodia: human macrophage-like THP-1, mouse macrophage-like RAW264.7, MDA-MB-231 human breast cancer cells, and MTC (mouse musculus thyroid carcinoma) cells. Podosomes in RAW264.7 and THP-1 cells were identified with imaging of actin core and/or vinculin ring. The latter two types of cells form invadopodia, which were identified with staining of actin and cortactin (Fig. S3). SNS assay was also applied to CHO-K1 cells as the negative control, which does not form podosomes or invadopodia. As expected, CHO-K1 cells did not form F-actin in a punctate pattern and exhibited no DNase activity on the SNS surface (Fig. 2a). In contrast, all the four invadosome-forming cell lines exhibit significant DNase activity localized with the actin cores of invadosomes (Fig. 2b-e), suggesting that DNase recruitment by invadosomes is highly consistent and ubiquitous in the different cell types and species. A statistical analysis shows that over 80% of podosomes in RAW264.7 and THP-1 have exhibited DNase activities. Percentage of invadopodia exhibiting DNase activities are about 80% in MDA-MB-231 cells and 46% in MTC cells (Fig. 2f).

**Fig. 2.**
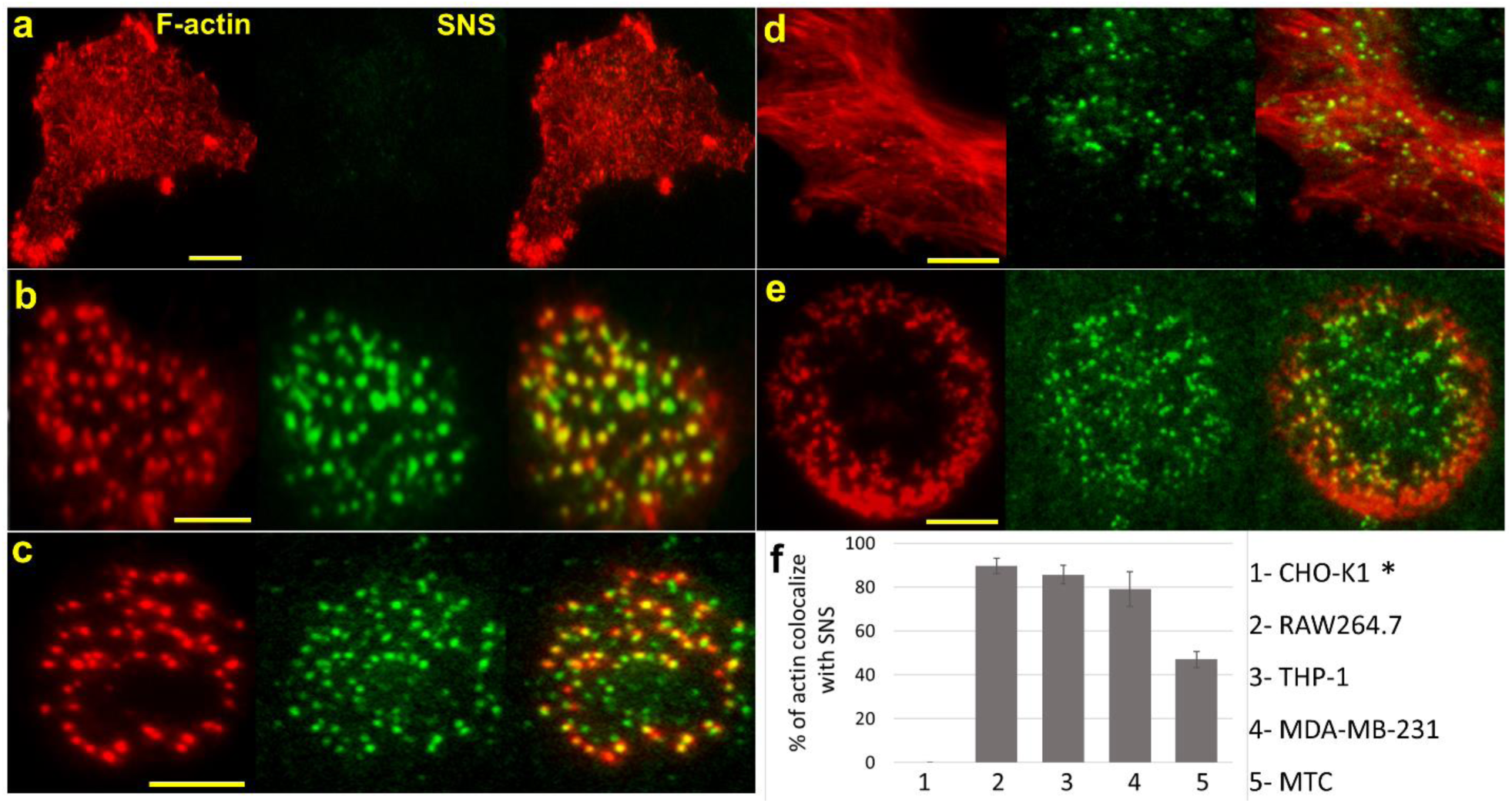
Ubiquitous DNase activity in invadosome-forming cells. (a) CHO-K1 cell does not form invadosomes (no actin core), and no DNase activity was observed at the cell-substrate interface. Strong DNase activity was consistently observed in a variety of cells that form podosomes or invadopodia. (b) RAW264.7 (c) THP-1 (d) MDA-MB-231 and (e) MTC, with former three forming podosomes and latter two forming invadopodia. Scale bar: 5 µm. (f) Percentage of actin cores that are co-localized with strong SNS punctate signal (20 cells and 400-700 invadosomes are analyzed for each cell type). *CHO-K1 does not form invadosome hence the colocalization was considered as zero.

### Temporal dynamics of DNase activity in podosomes

Podosome and invadopodia have different temporal dynamics in terms of their structure and function. To investigate the temporal dynamics of the DNase activity relative to the structural formation of podosomes, we tested RAW264.7 cells stably transfected with lifeAct-GFP that reports F-actin on an SNS surface (Fig. 3a, Video 1). Fluorescence time-lapse imaging with live cells was performed to monitor both the SNS signal and actin nucleation using a TIRF microscope. The result revealed that the DNase activity in podosomes manifested quickly (within ∼1 minute) after the initiation of actin nucleation and remains active (Figs. 3b, c) afterward, demonstrating strong enzymatic activity degrading extracellular DNA by podosomes. Further experiments were performed to test the correlation between the actin nucleation and DNase recruitment. It was found that the actin core formation is required for DNase recruitment to podosomes, as inhibiting actin polymerization with cytochalasin D abolished DNase activity in the punctate pattern, while inhibiting myosin II (another important structural component) with blebbistatin in podosomes had an insignificant effect on DNase activity (Fig. S4).

**Fig. 3.**
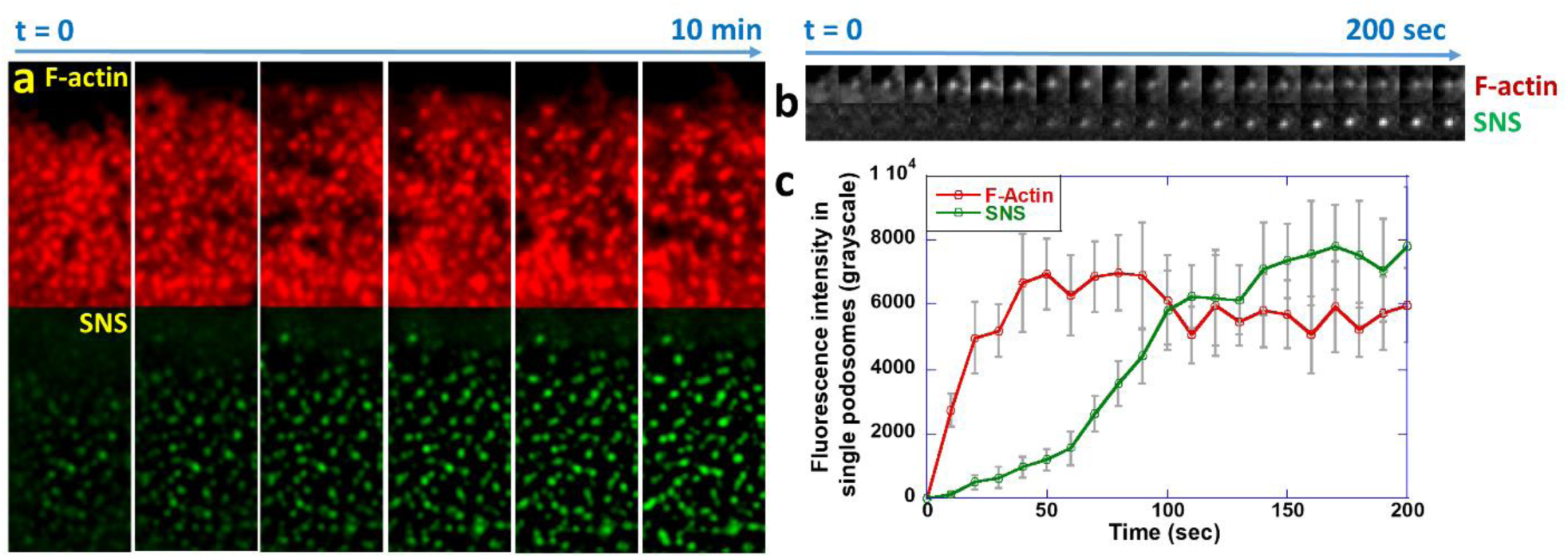
Temporal dynamics of actin core nucleation and DNase activity. (a) F-actin and SNS signal reporting DNase activity was co-imaged in RAW264.7 transfected with lifeAct-GFP. (b) Time-series images of F-actin and SNS signal in one example podosome (Video 1). (c) Temporal curves of F-actin and DNase activity averaged over 20 podosomes, showing that DNase activity manifests within about 1 minute after actin core nucleation.

### DNase in invadosomes is GPI-anchored DNase X

The localized DNase activity in a punctate pattern suggests that the DNase in invadosomes is likely a membrane-bound enzyme. Otherwise, the SNS signal would not be narrowly localized in invadosome because of the diffusion of soluble DNase. Motivated by this hypothesis, we first tested whether the DNase in invadosomes is membrane-anchored by GPI, which is a common lipid anchor linking a protein to the cell membrane. Treated with phosphatidylinositol-specific phospholipase C (PI-PLC), which cleaves the GPI anchor, DNase activity in podosomes in THP-1 cells was significantly reduced, and SNS signal was nearly zero under the treatment of 5 unit/mL PI-PLC (Figs. 4a, b). This result suggests that the DNase activity in podosomes is due to GPI-anchored DNase. The unaltered actin core structures in PI-PLC treated cells suggests that the reagent does not disrupt the structural formation of the podosomes. As DNase X was previously known as GPI-anchored membrane enzyme^31^, we further tested by immunostaining if the DNase in podosomes is DNase X. The immunostained spots of DNase X and the SNS puncta are well co-localized (Fig. 4c), confirming that the DNase in podosomes is DNase X. Collectively, these two experiments prove the presence of GPI-anchored DNase X specifically at podosome sites in THP-1 cells.

**Fig. 4.**
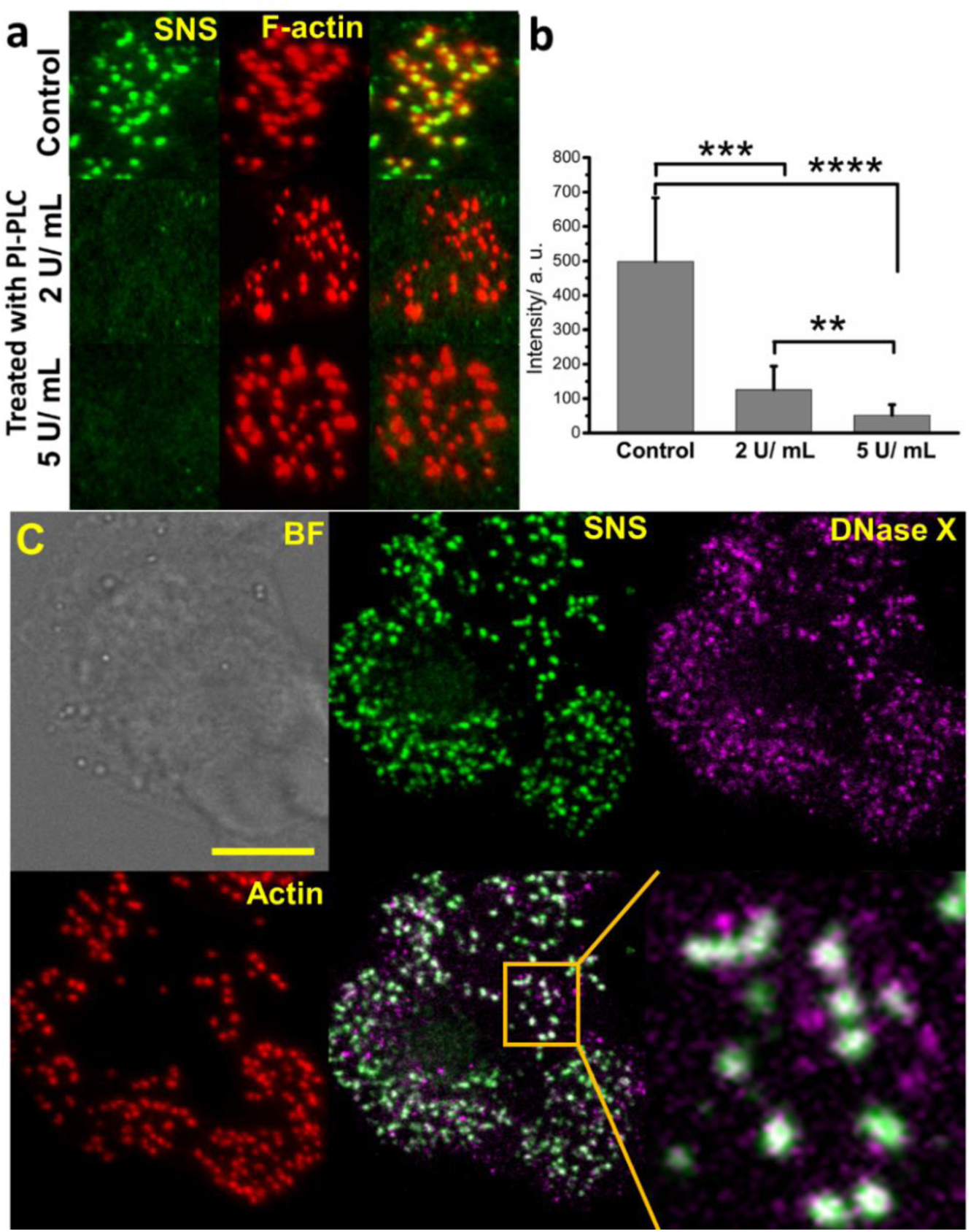
Identification of DNase active in invadosomes. (a) SNS and F-actin signals in THP-1 cells treated with PI-PLC, which cleaves GPI anchors. (b) SNS signals in THP-1 cells based on 20 cells, showing that PI-PLC significantly reduced the DNase activity in invadosomes. (c) Immunostaining of THP-l cells with anti-DNase X showed good colocalization between anti-DNase X and SNS signal. Scale bar: 10 µm.

### Capability of degrading DNA and matrix proteins by invadosomes

Enzymatic degradation is the crucial biochemical function of invadosomes. Previously, extensive research has been dedicated to investigating the degradation of ECM proteins by invadosomes^29, 32^. However, ECM also consists of extracellular DNA originating from apoptotic and necrotic cells, the NET (neutrophil extracellular traps)^33^, invading pathogens, *etc*.^33-34^ Here we designed a co-digestion assay to simultaneously monitor the degradation of both matrix protein and extracellular DNA by invadosomes. We coated glass surfaces with both SNS and dye-labeled (HyLight488) fibronectin (FN) and tested THP-1 cells and MTC cells on these surfaces. Time series imaging showed that podosomes were formed in activated THP-1 cells and rapidly degraded both SNS (made of dsDNA) and FN nearly simultaneously (Figs. 5a, b and Video 2). This suggests that DNase may potentially participate in matrix degradation, with an enzymatic role parallel to proteases. In contrast, invadopodia formed in MTC cells exhibited rapid DNase activity, but no degradation of FN due to protease activity was observed within our time frame of imaging (1 h) (Fig. 5c, d, Video 3). Some previous studies showed that invadopodia exhibited detectable protease activity on the gelatin surface within 1 h^35^. This discrepancy may be due to the fact that our assay was performed on the hard glass surface, where invadopodia may have longer maturation time than those on the soft gel surface. Nonetheless, our results suggest that DNase activity is more prompt and ubiquitous than protease activity in invadosomes, potentially making it a superior biomarker for the identification of invadosomes.

**Fig. 5.**
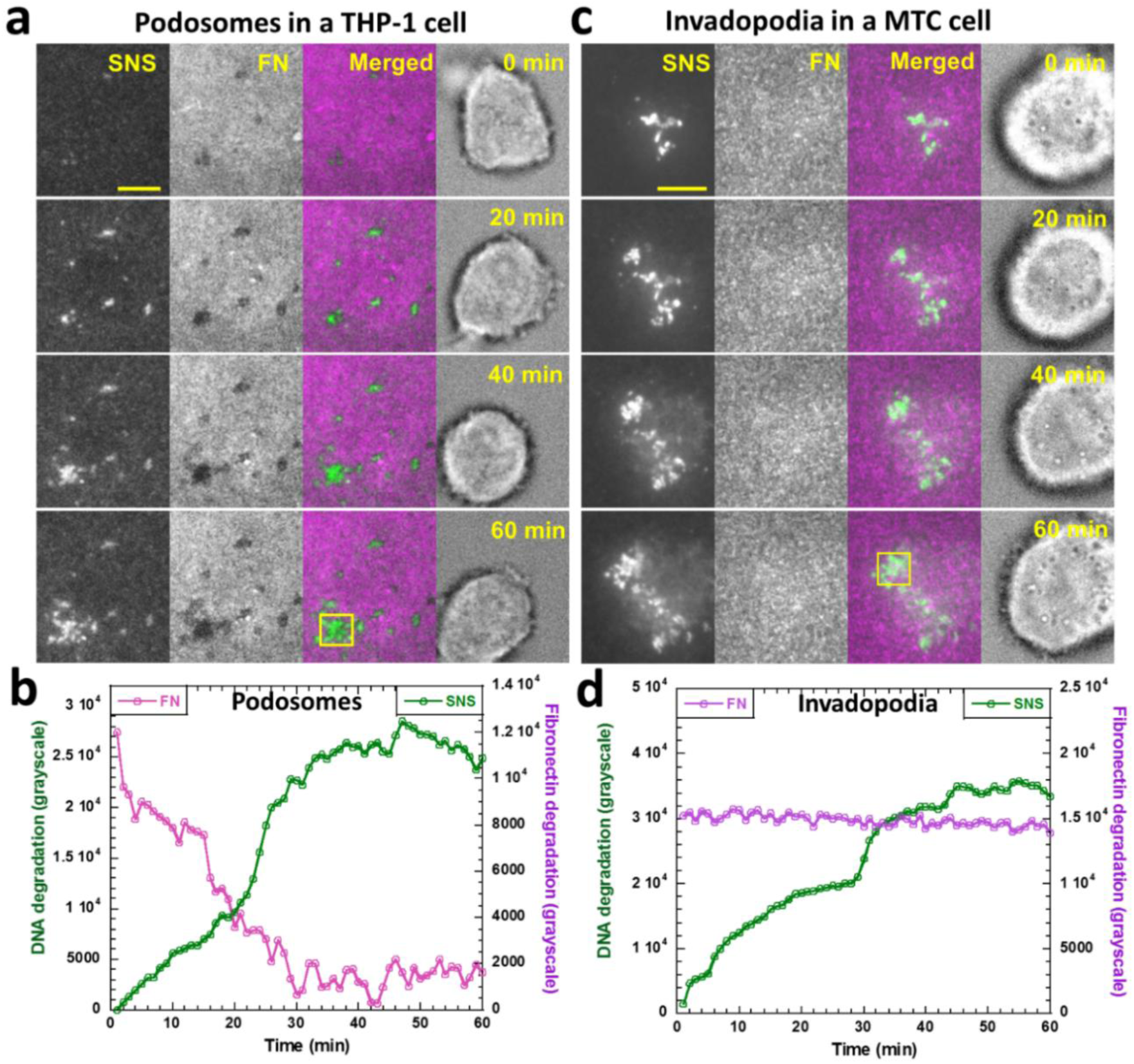
Co-degradation of SNS (extracellular DNA) and fibronectin (matrix protein) by invadosomes. (a) A podosome forming THP-1 cell was imaged over 60 min on a surface coated with both SNS and HyLite488-labeled FN (Video 2). (b) Fluorescence intensity analysis shows that the degradation of both SNS and FN is starting at a similar time by podosome. (c) Invadopodia forming MTC cell on a surface coated with both SNS and FN (Video 3). (d) Fluorescence intensity analysis shows that only SNS was degraded by invadopodia within the time frame of the experiment. Scale bar: 5 µm.

## Conclusions

In summary, we reported the ubiquitous DNase activity in both podosomes and invadopodia, characterized its temporal dynamics, and confirmed its identity. The DNase activity in invadosomes is strong, prompt, localized, and consistent in invadosomes across different cell types and species, suggesting that invadosomes have an enzymatic arsenal richer than previously known. The DNase may work with proteases in parallel to degrade ECM, which consists of both matrix proteins and extracellular DNA released by dead cells (*e.g.* neutrophil extracellular traps) and pathogens^33, 36^. DNase activity is more prompt and consistent than that of protease in invadosomes, potentially making it a more robust bio-marker of invadosomes to set them apart from other cell-adhesion units such as focal adhesions, lamellipodia, and filopodia^32^. The presence of DNase in invadosomes also provides new therapeutic targets and cautions for researchers in performing gene-based therapeutics to invadopodium-forming cancer cells.

Overall, revealing DNase in invadosomes adds an important piece in the whole picture of understanding the enzymatic behaviors of these dynamic and versatile structures. To the best of our knowledge, this is the first report of DNase activity in invadosomes since their discoveries decades ago. The absence of any study related to DNase in invadosomes in the past is likely because of the lack of a sensor that can image the DNase activity at the cell-substrate interface. This work will likely intrigue more research to study the physiological roles of DNase in invadosomes and the associated signaling pathways, and the DNase sensor SNS would be a valuable tool to bring the DNase to light.

## Materials and Methods

### Synthesis of SNS

We customized and ordered oligonucleotides from Integrated DNA Technologies (IDT) with sequences and modifications as shown below (Sequence from 5′ end to 3′ end):

DNA 1: GGG CGG CGA CCT CAG CAT/3BHQ_2/

DNA 2: /5BiosG/T/iCy3/ATG CTG AGG TCG CCG CCC/

DNA 3: GGG CGG CGA CCT CAG CAT

DNA 4: /5Cy3/ AGG TCG ATG CTG CCG CCC/3Bio/

For the construction of SNS, DNA 1 and DNA 2 dissolved in 1× phosphate buffered saline (PBS) were annealed at a molar ratio of 1.2:1.0 and stored at 4 °C at a final concentration 10 µM for further use. For the construction of Cy3-labeled DNA, DNA 3 and DNA 4 were annealed in the same conditions as SNS annealing.

### SNS immobilization on the glass surface

The imaging platform for our study is the SNS-immobilized glass surface. To prepare this surface, we follow these steps. Step 1: a solution of 200 *μ*g/mL BSA-biotin (biotin conjugated Bovine serum albumin, A8549, Sigma-Aldrich, USA) and 2 *μ*g/mL fibronectin (1918-FN, R&D System, USA) in PBS buffer was incubated on a glass-bottom petri dish (D35-14-1.5-N, in vitro Scientific, CA, USA) for 30 min at 4 °C. BSA and fibronectin both can be physically adsorbed on the glass surface. Fibronectin was used for assisting cells to adhere. Then the surface was rinsed with cold PBS thrice. Step-2: Previous surface was incubated with a solution of 50 μg/mL neutravidin (31000, Thermo Fisher Scientific, MA, USA) in PBS for 30 min at 4 °C and rinsed with cold PBS thrice. Step-3: Previous glass surface was incubated with the solution of 0.1 *μ*M SNS in PBS for 30 min at 4 °C and rinsed with PBS thrice. The glass surface became ready for cell plating and further experiments.

### Fibronectin-HiLyte 488 surface preparation

A solution of 5 *μ*g/mL Fibronectin labeled with HiLyte fluor 488 (FNR02-A, Cytoskeleton, Inc. USA) in PBS was incubated on glass bottom petri dish for 30 min at 4°C and rinsed with cold PBS thrice. The petridish was further coated with 200 *μ*g/mL BSA-biotin, 50 *μ*g/mL neutravidin and 0.1 μM SNS, consecutively, and became ready for experiments.

### Cell culture and cell plating

Both cancer (MDA-MB-231 and MTC) and non-cancer macrophage resembling (RAW264.7 and THP-1) cell-lines were used in SNS assays. All the cell lines were cultured according to the suppliers’ protocol. The macrophage activation was done with the treatment of 1 ng/mL lipopolysaccharide (L2630, Sigma Aldrich, USA) to RAW264.7 or 2 ng/mL transforming growth factor *β*-1(240-B/CF, R&D System, USA) to THP-1 for 24-36 hours, respectively. Here the RAW264.7 cells were stably transfected with LifAct-GFP. For experiments, cells were harvested at 80–90% confluency using the following method: After removing the culture medium, cells were firstly washed and incubated with cell detaching solution inside the CO2 incubator for 5-7 min. [recipes for 1L cell detaching solution: 100 mL 10×HBSS +10 mL 1 M HEPES + 10 mL 7.5% sodium bicarbonate +2.4 mL 500 mM EDTA (Ethylenediaminetetraacetic acid). Rest of the volume was made up with distilled water and PH was adjusted to 7.4]. Cells were pipetted off gently from the petri dish and transferred into a 1.5 mL centrifuge tube and centrifuged for 3 min at 300g. Cell pellet was re-suspended with the corresponding complete medium. The solution was then transferred on the glass surface modified with SNS and incubated for 30-60 min inside CO2 incubator before imaging.

### Immunostaining

Immunostaining was started with the fixation of the cells with 4 % formaldehyde solution in PBS for 15 min at room temperature. After removing the formaldehyde solution, cells were permealized with 0.1% Triton X-100 solution in PBS for 20 min at room temperature. Then the cells were blocked with 5 mg/ mL BSA solution in PBS for 45 min at room temperature. For the staining of actin, Phalloidin-Alexa 405 (A30104, Thermo Fisher Scientific, USA) was used and the sample was incubated according to the supplier’s instruction and washed with PBS for three times with 5 min each. For the staining of vinculin, tumor necrosis factor *α* (TNF*α*) or DNase X, cell samples were first treated with primary antibodies: anti-vinculin (90227, Millipore), anti-TNF*α* (ab1793, abcam, UK) or anti-DNase X (H00001774-MO2, Abnova, Taiwan) in PBS for 1h at room temperature, and then washed with PBS for three times. PBS remained on the samples for 5 min prior to its removal. The samples were then treated with dye-labeled secondary antibody (ab175660, abcam, UK) solution in PBS and incubated for 1h at room temperature. Wash thrice with PBS with 5 min incubation time. The cortactin staining was performed with dye-labeled Anti-cortactin (05-180-AF647, Millipore) and therefore the secondary antibody treatment was skipped.

### Cell transfection

For the transfection, corresponding cells were cultured in a 35 mm petri dish up to 70-80% of confluency. Plasmid DNA (2-3 *μ*g) was diluted in 400 *μ*L Opti-MEM (11058021, Thermo Fisher Scientific, USA) spiked with 2-3 *μ*L Plus reagent (15338100, Thermo Fisher Scientific, USA). The plasmid solution was mixed and incubated for 10 min at room temperature. After that, 6-8 *μ*L Lipofectamine-LTX (15338100, Thermo Fisher Scientific, USA) was added to the plasmid solution which was then mixed and incubated at room temperature for 30 min. The medium in petri dish of cells was exchanged with 2 mL fresh complete medium added with the plasmid mixture. Cells were incubated in an incubator for 18-24 h. Before experiments, cells were detached with EDTA solution and plated on the imaging platform.

### Inhibitor study

GPI inhibition: THP-1 cells activated by 2 ng/mL TGF-*β*1 (Transforming growth factor *β*1) for 36-48 h were plated on the SNS imaging plate along with 2 or 5 unit/mL of PI-PLC (P8804, Sigma Aldrich, USA) in complete growth medium for 1 h and then stained with phalloidin-Alexa 405 prior to imaging. Other Inhibitor: activated cells were detached from the surface with EDTA treatment and then then centrifuge them in a centrifuge tube. Now the pallet is resuspended in complete medium with desired concentration of inhibitor, namely blebbistatin (Sigma Aldrich, B0560) or cytochalasin D (Sigma Aldrich, C8273). Now this cell suspension with inhibitor was plated on the imaging substrate and incubated in CO2-incubator before imaging.

### Microscopy and image processing

All static and time-lapse microscopy imaging was performed using total internal reflection (TIRF) microscopy setup (Nikon Ti-2) with a 100X oil immersion objective. Images and Data were analyzed by Matlab code developed in our lab.

## Acknowledgements

We thanks Dr. Kaitlin Bratlie for sharing the RAW264.7 cell-line and Dr. Ian Schneider for sharing MTC cell-line.

## Competing interests

The authors declare that they have no competing interests.

## Notes

### Competing Interest Statement

The authors have declared no competing interest.

