## Supplemental text for "Fluorogenic DNase Sensor Reveals Ubiquitous DNase Activity in Podosomes and Invadopodia"

### Video Captions

Video 1. Co-imaging of F-actin and DNase activity in a RAW264.7 cell on a SNS surface.

Video 2. A THP-1 cell degrades DNA and fibronectin.

Video 3. A MTC cell degrades DNA but not fibronectin within the experiment time frame.

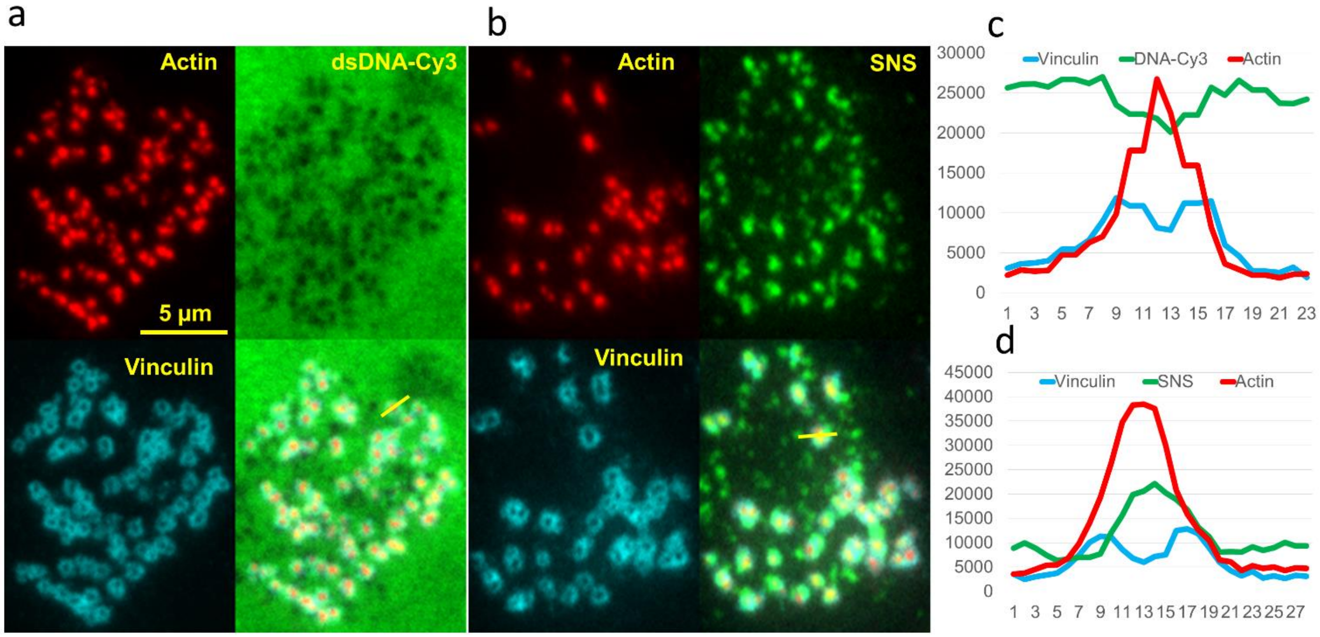


**Fig. S1.** Co-localization analysis of DNase activities and structural proteins in podosomes. A THP-1 cell on Cy3-labeled dsDNA (a) and SNS (b) modified surface showing fluorescence loss and gain due to DNA degradation by the DNase in podosomes, which are identified with actin cores and vinculin rings. The DNase activity is well situated in the core region as the dark (c), and bright (d) punctate signal is co-localized with actin core and void area of vinculin ring. The intensity is plotted along the solid yellow line.

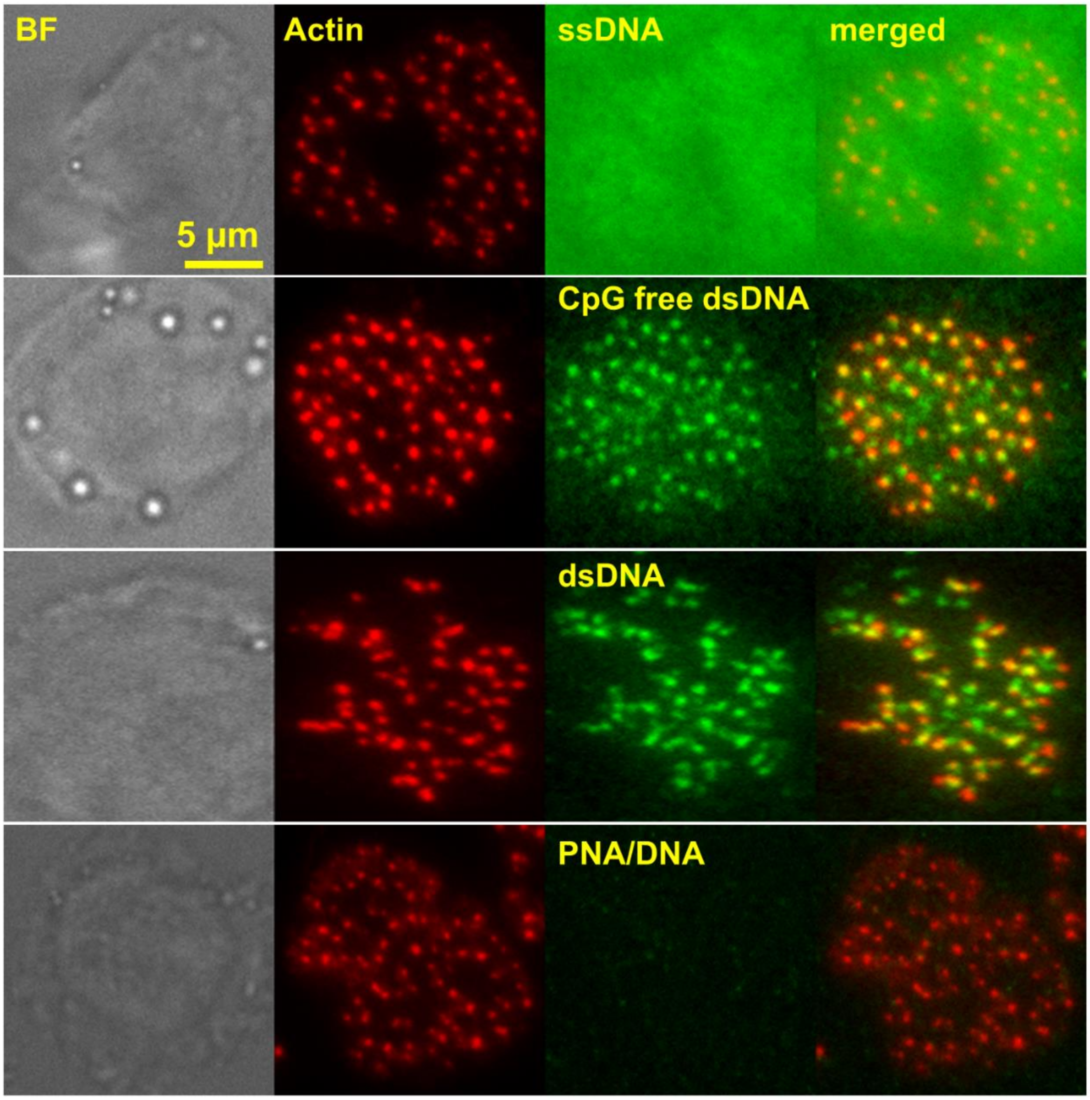


**Fig. S2.** DNase in podosomes does not digest ssDNA but degrade dsDNA irrespective of their sequences. THP-1 cells were plated on ssDNA, CpG cluster free dsDNA (CpG motif potentially stimulates immune response), dsDNA and DNA/PNA modified surface where podosomes (identified with actin core) degraded the dsDNA regardless of DNA sequences but no enzymatic effect was exhibited on ssDNA or PNA/DNA hybrid.

All DNA sequences are from 5' end to 3' end:

**ssDNA**: 5Biosg/T/iCy3/ATG CTG AGG TCG CCG CCC

**CpG free**: GGCACCCAGGCACCACCC/3BHQ2
5Biosg/T/iCy3/GGGTGGTGCCTGGGTGCC

**Other Seq**: GGGAGGACGGAGCAGGGC/3BHQ2
5Biosg/T/iCy3/GCCCTGCTCCGTCCTCCC

**PNA/DNA**: GGGCGGCGACCTCAGCAT/BHQ2
Biotin-OO-Lys(Cy3)-O-ATGCTGAGGTCGCCGCCC

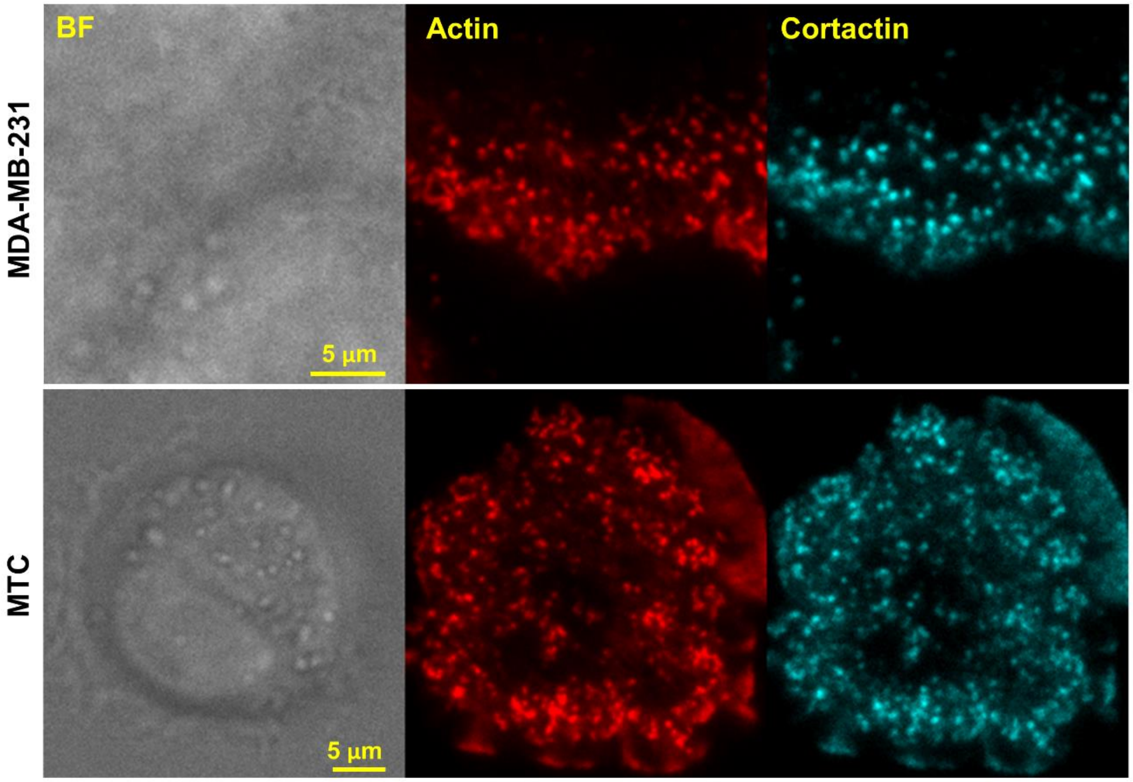


**Fig. S3.** Characterizing actin puncta as the invadopodia in two cancer cell lines. Cancer cell line MTC and MDA-MB-231 had a number of punctate actin structures. These actin puncta were identified as invadopodia as they were well co-localized with cortactin shown by immunostaining.

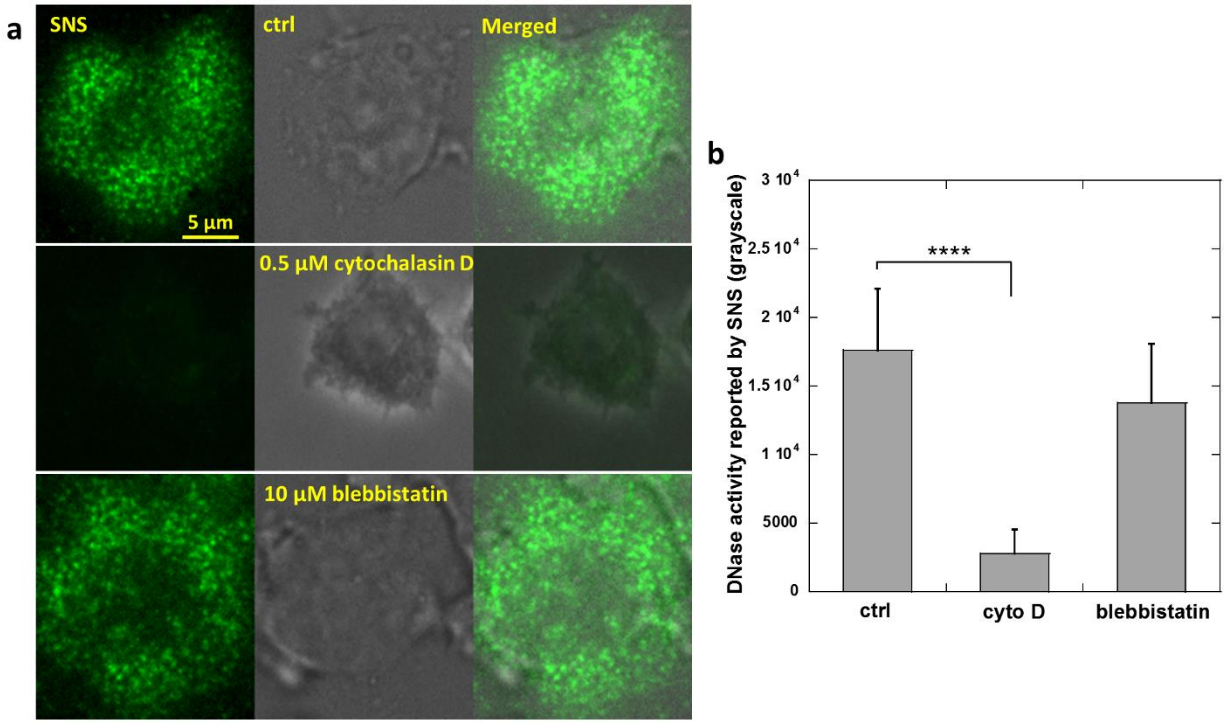


**Fig. S4,** Actin polymerization is required for DNase activity in podosomes. (a) RAW264.7 cells were plated on SNS surfaces with actin polymerization inhibited by cytochalasin D or myosin II inhibited by blebbistatin. (b) The bar graph showed that the cytochalasin D significantly reduced the DNase activity in podosomes whereas blebbistatin had a minor effect on DNase activity.
